# G-Trap Assay II: Characterization of blood Leukocyte Functionality differentiates immune activation and immune suppression in bacteremia patient samples

**DOI:** 10.1101/2022.06.09.495553

**Authors:** Peter Simons, Laura Shevy, Virginie Bondu, Angela Wandinger-Ness, Stephen Young, Tione Buranda

## Abstract

Sepsis is a severe organ dysfunction syndrome caused by a dysregulation of the immune system’s response to infection. Unfortunately, most infection-causing pathogens aren’t routinely detectable in real-time to enable targeted and lifesaving treatment. Thus, clinicians frequently have limited data on which to base treatment decisions. A complete blood count with differential is available within 24 h, and positive culture is only available in ~30% of cases. Furthermore, a blood culture, the traditional gold standard for accurate diagnosis of bacteremia, may take up to five days for results, long after a clinical decision for sepsis management is required. Circulating leukocytes can sense chemotactic signals released by bloodborne pathogens or focal infections not in the bloodstream. Our earlier study showed that pathogen and host immune factors released in the bloodstream stimulated GTP binding of Ras homology (Rho) GTPases (guanosine triphosphatase) such as Rac1 in quiescent endothelial and human leukocytes after exposure to blood plasma from infected patients.[1] In this study, we measured Rac1•GTP as a biomarker of immune functionality of peripheral blood monocytes and polymorphonuclear cells extracted from blood samples drawn for diagnostic use in blood culture assays; from 120 non-infected control patients and serial blood test samples from 28 patients with a confirmed diagnosis of bloodstream infection. 18 cases presented with Rac1•GTP elevation of ≥3 fold above that of control samples. Ten patients with normal or below-normal GTPase activity, accompanied by neutrophilia or pancytopenia. We used Principal Component Analysis to differentiate the 2D spatial distribution of infected patients and negative controls. Measuring differential leukocyte functionality in infected and control patients’ blood samples with the G-Trap assay may provide an innovative process for a real-time distinction between infection and non-infectious etiologies.

## INTRODUCTION

Sepsis is a life-threatening organ dysfunction syndrome caused by a dysregulated host response to infection. Bacterial bloodstream infections (bacteremia) are the most common precursor to sepsis. Blood culture is the most common diagnostic test used to confirm bacterial sepsis. However, most acute bacterial infections are tissue localized and isolated from the bloodstream. Additionally, pathogen-focused methods can take hours or days to return a result. The sensitivity and specificity of blood culture are low; 30% positivity due to several factors such as limiting blood volume drawn, a temporal mismatch between the blood draw time and bacteremia onset, prophylactic or empiric antibiotic treatment, and the presence of viable organisms. To narrow antimicrobial therapy selection, blood culture-adjacent assays such as genotypic or phenotypic antimicrobial susceptibility testing (AST)[2-5] can be used for rapid post-blood-culture bacterial PCR panels day. More recently, targeting the host response has emerged as potentially the best way to enhance initial sepsis workflows for diagnosing and treating sepsis. Conventional markers include white blood cell (WBC) counts, C-reactive protein, and procalcitonin. Procalcitonin is most specific as a longitudinal marker to determine antibiotic efficacy.[6, 7] However, its use has not produced a significant change or improvement in antibiotic stewardship.[8, 9] Host immune response mRNA panels integrated with machine learning algorithms to accurately identify the presence and type of infection (virus, bacteria, fungus) anywhere in the body have been recently described. [9-12]

Sepsis may cause either leukocytosis or leukopenia, with other bacteremic patients presenting to the hospital with a normal WBC.[13] Thus, in some respects, WBC count has limited utility as a tool for infection diagnosis. [14] Sepsis often manifests as a complex immune dysfunction syndrome with a simultaneous presentation of inflammatory and anti-inflammatory responses.[15, 16] It is worth noting that sepsis and cancer present similar immune dysfunction in innate and adaptive immune cells.[17] Under sepsis, immune cells’ life span, production, and function are susceptible to modification, often linked to anergy.[18-20] During the acute phase of the disease, the immune response is characterized by inflammation and the activation of immunological effector mechanisms. Damaged cells and tissue-resident macrophages that have ingested pathogens release proinflammatory cytokines and chemotactic cytokines[21,22] to recruit effector innate immune cells that lack antigen specificity (such as neutrophils, monocytes, and macrophages) and adaptive immune effectors (such as T and B memory lymphocytes). [23-30]

Rho and Ras family GTPases are critical effectors of chemoattractant-mediated motility of effector cells towards sites of infection and pathogen killing (**Fig. 1**). [31-37] The GTPases function as molecular switches, cycling between inactive GDP-bound and active GTP-bound states. GTP-bound proteins specifically bind to effector proteins to advance GTPase biological function. [38] Several studies have shown that impaired activation of adaptive immune cells is associated with failure to induce GTP loading to leukocytes.[34, 39-42] After clinical recovery, sepsis might confer long-term deficits (immune suppression, chronic inflammation) of innate and adaptive immune function and serial persistence of nosocomial polymicrobial infections.[43, 44] Monitoring immune function for sepsis-associated immune cell defects could be used to improve the potential for infection diagnosis and immune modulatory therapy to improve long-term patient outcomes.

**Fig. 1.**
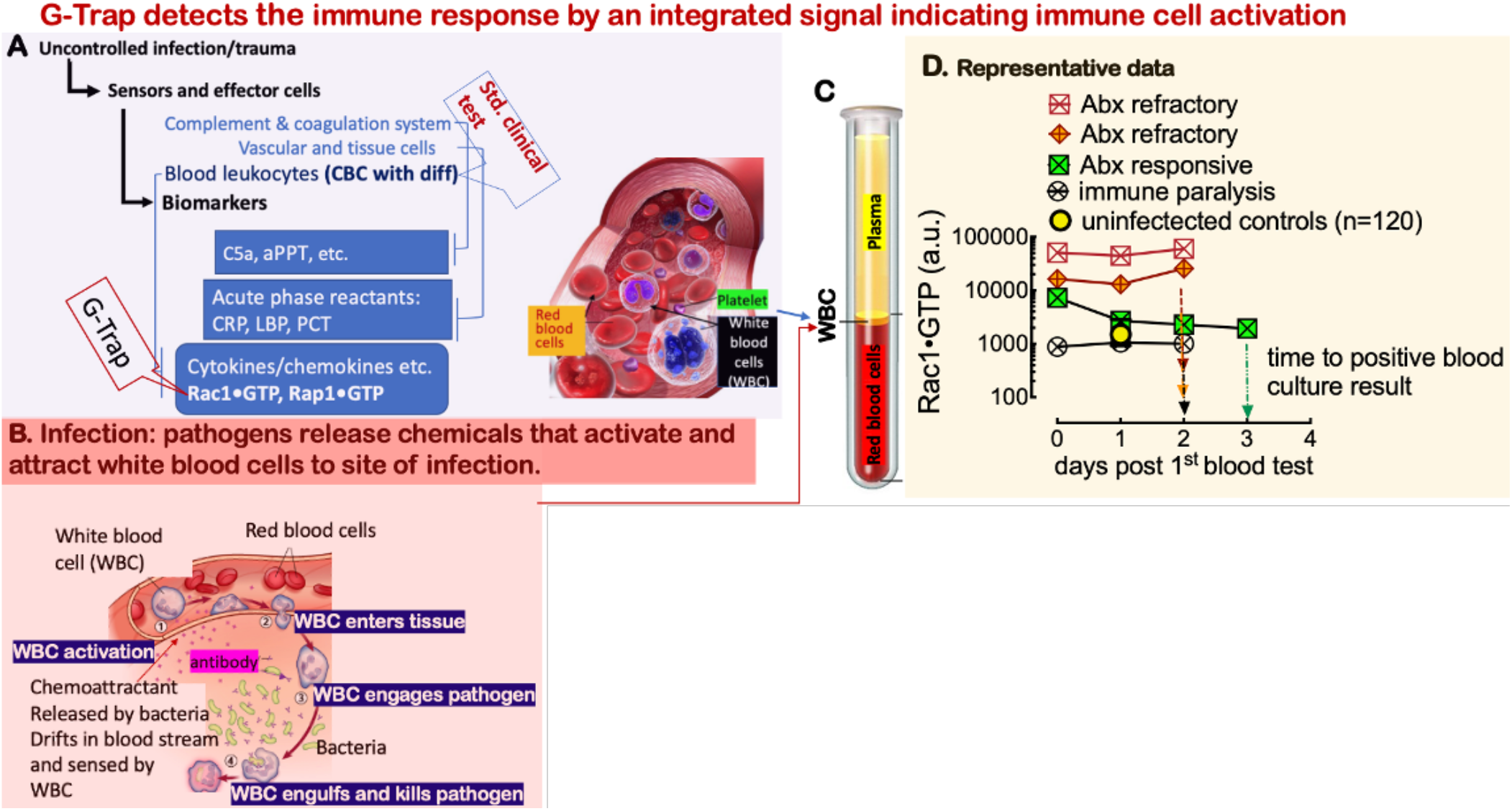
**A**. Bloodstream bacterial infection is detected by host innate immune cells, including neutrophils, monocytes, and dendritic cells, which activate adaptive lymphocyte effector functions. Changes in leukocyte counts due to infection are routinely measured by a complete blood count with differential (CBC with diff). However, various immune response biomarkers have proven too variable to be definitive diagnostically. The G-Trap assay measures levels of Rac1•GTP, among other activated GTPases, in lysates of circulating white blood cells (WBC). Active GTPases serve as sensors for infection due to their roles in triggering WBC to migrate to sites of infection and kill pathogens. **B.** Bloodstream bacterial infection is detected by white blood cells. Active GTPases serve as sensors for infection due to their roles in triggering WBC to migrate to sites of infection and kill pathogens. Changes in white blood cell (WBC) counts due to infection are routinely measured as an indicator of infection. (image adapted from https://together.stjude.org/en-us/care-support/immunity-illness-infection.html) **C.** Separation of whole blood components allows isolation of WBC, which are then analyzed using the G-Trap assay. Image of Blood cells retrieved on April 15, 2022, from (https://www.visiblebody.com/learn/biology/blood-cells/blood-overview) **D**. Rac1•GTP is a sensitive biomarker of pathogen-activated white blood cells. Serial analysis of blood samples of representative patients presenting an active immune response: two patients showed a failure of antibiotic treatment one patient was responsive to treatment. One patient indicated immune paralysis indicated by below normal GTPase activity.

In a preceding study, we used serial plasma samples from infected patients to simulate immune activation of human leukocytes and culture cells after exposure to inflammatory mediators released in the blood circulation of infected patients. That study showed that pro-/anti-inflammatory cytokines and novel biomarkers such as chemerin[45] induced GTPase activation in test cells. In this study, we hypothesized that Rho GTPase activation in response to infection[31] is required for immune effector function, such as chemotaxis and tissue extravasation of circulating blood leukocytes during the early and late stages of an infection.[23-30] **(Fig. 1A-C**). Immune activation and dysfunction highlight the need for an assay to assess the functionality of white blood cells, which are an essential component of diagnosing the potential of an infection. The small GTPase Rac1 plays a primary role in cell motility.[46] Herein we measure GTP loading to Rac1 in peripheral blood mononuclear cells (PBMCs) and polymorphonuclear cells (PMNs) isolated from controls and serial clinical diagnostic samples used to confirm bacterial infection from blood culture. Our data indicate that the GTPase activity as a measure of functionality of PBMCs and PMNs is a promising biomarker of immune cell function and could potentially be used as a clinical decision support tool (**Fig. 1D**).

## RESULTS AND DISCUSSION

### GTP binding to Rac1 in patient PBMCs is sensitive to environmental allergies, whereas GTP binding to Rac1 in PMNs is not

We tested peripheral blood samples from 120 non-infected subjects to establish a baseline assessment of GTPase activity in uninfected control patients. The samples were collected over 6 months, from August to December 2019. On average, 12 unique patient samples were tested each week. During the August-October period, we noted that the median level of GTPase activity for PBMCs was variable each week, whereas the GTPase activity of PMN was stable (**Fig. 2A&B**). Therefore, we hypothesized that changes in local pollen counts drove T_h_2 cell-mediated allergic response[47, 48], which caused the variation in GTP binding to Rac1 in PBMCs. Consistent with our hypothesis, a plot of the pollen counts (Weather.com) in Albuquerque on the days the blood samples were collected showed a strong and significant Pearson correlation (ρ = 0.78, p = 0.005) between the median GTPase activity of PBMC samples and pollen counts. We, therefore, included this signal variability within the signal range expected of control samples.

**Fig. 2.**
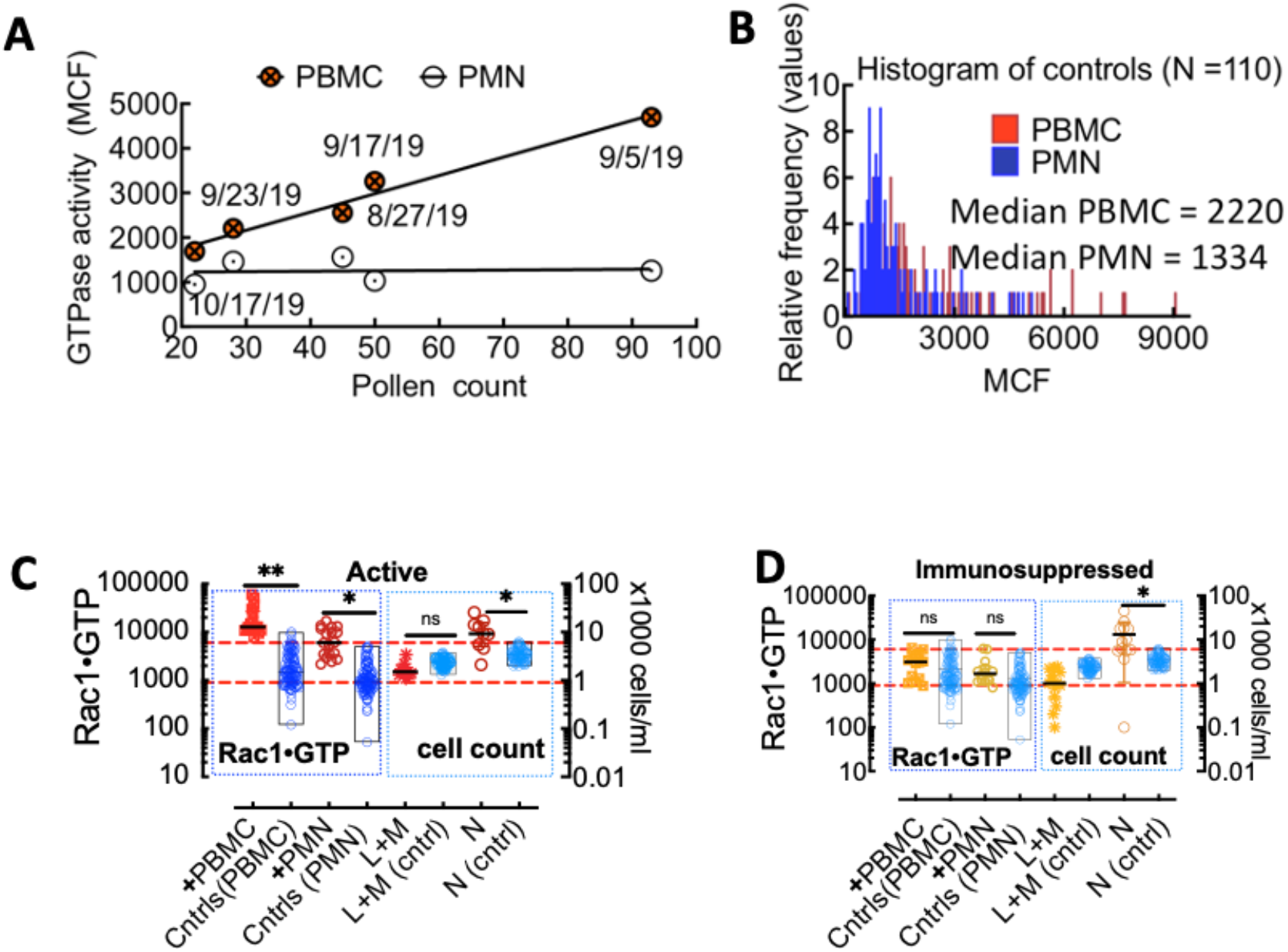
**A.** Correlated changes in GTPase activity in PBMCs and pollen count on the day of blood sample collection. Each day represents 12 samples. Pearson correlation (ρ = 0.78, p = 0.05). **B.** Overlapping histograms of Rac-1 GTPase activity measured with G-Trap in leukocytes from TriCore diagnostic blood draws of uninfected patients. The fluorescence value for each patient is represented as a Median Chanel fluorescence (MCF) of the G-Trap measurement. The patient population’s median values for polymorphonuclear neutrophils (PMN) and peripheral blood mononuclear cells (PBMC) are 1334 and 2220, respectively. **C.** Plot of Rac1•GTP measured in cell lysates of normal infection-positive patient leukocytes (+PBMC and +PMN) and controls. The plot of Rac1•GTP was measured in cell lysates of infection-positive patient leukocytes (+PBMC and +PMN) and controls. **D.** Plot of Rac1•GTP measured in cell lysates of infection-positive immunosuppressed patient leukocytes (+PBMC and +PMN) and controls. Levels of Rac1•GTP were compared using unpaired t-test; ** *p*< 0.01, * *p*< 0.05.

### Rac1 • GTP levels are significantly upregulated in patients presenting a normal immune response to infection and are depressed in patients with apparent immunosuppression

We assayed for Rac1•GTP in serial blood samples drawn from patients with culture-positive bacterial infections from 28 patients. The peak GTPase activity in PBMCs from 18 out of 28 infected patients was ≥3-fold higher than the activity (interquartile range) of the control samples. Lymphocyte and monocyte counts in infected patients were lower than in uninfected controls. However, the difference was not significant. Sepsis profoundly affects all the main lymphocyte subpopulations [43, 49] which undergo increased apoptosis; T-regulatory cells are more resistant to sepsis-induced apoptosis, leading to an increased proportion of T-regulatory cells and an immunosuppressive phenotype. Neutrophil (N) counts were significantly elevated relative to controls (**Fig. 2C**). Overall, Peak Rac1•GTP was lower in PMNs than PBMCs regardless of the higher cell counts in the former.

Rac1•GTP measured in samples from 10 of 28 cases was comparable to controls or at the lower end of the mean quartile range, even though the patients’ neutrophil counts were significantly elevated compared to controls (**Fig. 2D**). We assessed the average leukocyte functionality for single cells in each sample population by dividing the Rac1•GTP expression of each patient sample by its cell count (**Fig 3**). The difference between infected and control Rac1•GTP/cell data was significant for PBMCs and not in PMN samples. This result suggested an inverse correlation between neutrophilia and functionality. A notable exception was patient #14, presenting with severe neutropenia (dy2 = 0.09, 1.4k/μl; Table S1), yet yielded a robust GTPase response. We found similar trends for patients # 24, 28, and 31 due to lowered cell counts. We plotted the temporal Rac1•GTP data associated with PBMCs and PMNs from representative patient samples (**Fig 4A-B**). Normalization of GTPase activity to single cells enhanced separation of immune activity levels among the patients according to differences in cell counts (**Fig. 4C-D**), also notably highlighting patient #14’s depleted but very active neutrophils (**Fig. 4D**).

**Figure 3.**
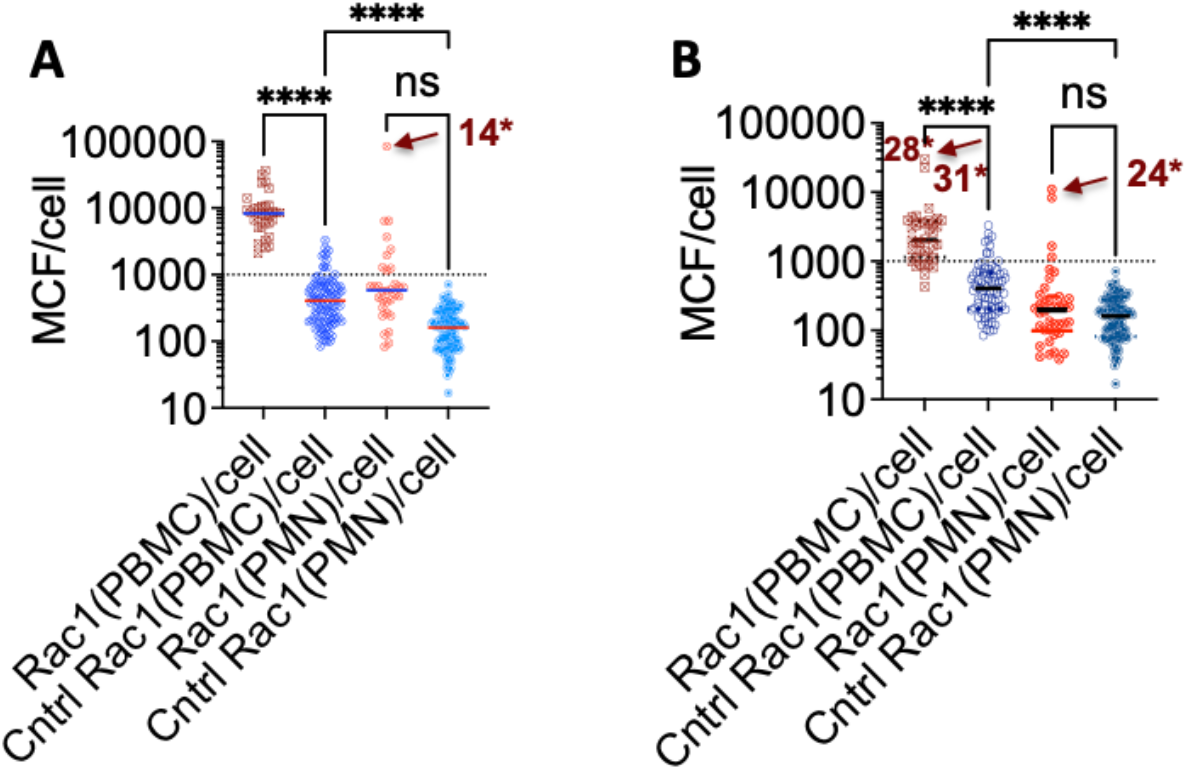
Immune functionality of PBMCs and PMNs per unit cell. **A.** Patient samples presenting GTPase activity above at ≥3 fold threshold above the median value of the control. **B**. Patient samples presenting GTPase activity at <3 fold median value of the control. The arrows indicate robustly active cellular immune responses of severely neutropenic (14 and 24) or lymphopenic (28 and 31) patients. Levels of Rac1 were compared by ordinary one-way ANOVA with Tukey’s correction for multiple comparison tests; **** *p*< 0.0001. Please refer to Supplementary Table 1 for raw data reference.

**Fig. 4.**
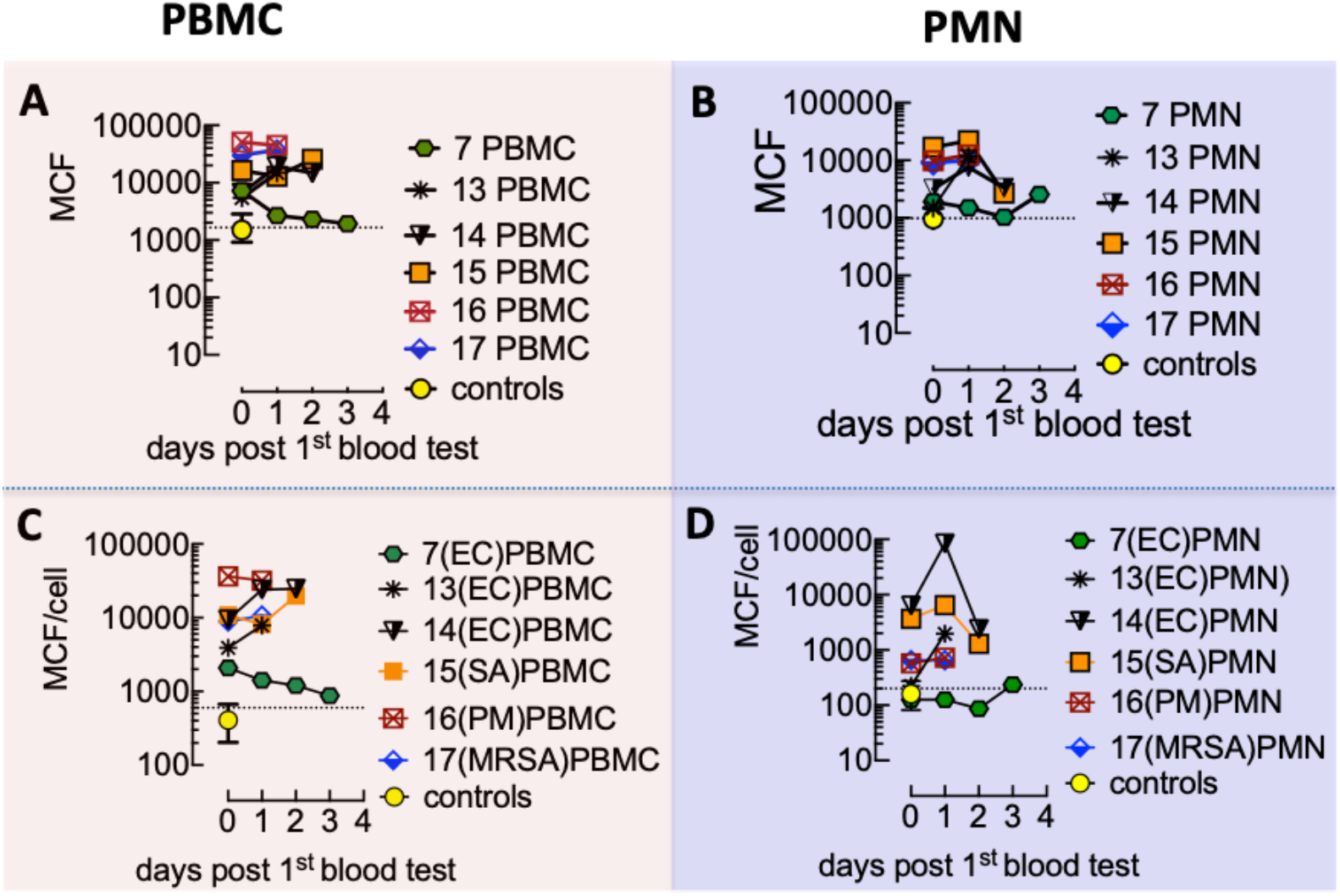
Representative graphs of Rac1•GTP measured in serial lysates of PBMCs and PMNs from patients presenting GTPase activity ≥3fold above controls. **A&B.** Plots of Rac1•GTP in measured in PBMC and PMN lysates versus sample-collection day post-admission and before positive culture result. Each patient is identified numerically. Infecting pathogen abbreviations in parentheses and (G+ or G-) refer to Gram staining of the organism: (PM) Pasteurella multocida (G-); (SA) Staphylococcus aureus (G+), (MRSA) methicillin-resistant Staphylococcus aureus (G+); (EC) Escherichia coli (G-).**C&D.** Rac1•GTP normalized to cell count for the same patient group.

We next used Principal Component Analysis (PCA) [50] to determine the relationship between the relative expression of different types of significant leukocyte groups: lymphocytes (L), monocytes (M), and neutrophils (N) and their immune function represented by Rac1•GTP measured in PBMCs and PMNs from patients with confirmed bloodstream infection compared to uninfected controls. (**Fig. 6**). The data are presented using Graphpad prism as a distance biplot of Principal component (PC) scores (patient samples) and *PC loadings* vectors (Rac1 (PBMC), Rac1(PMN), N, L, M, and L+M). The *PC loadings* vectors indicate a proportionate concentration of the immunological variables and correlation with serial patient samples. The coordinates of the vector end or patient samples indicate the weight of the immune variable and the relationship (correlation) to patient samples.

**Fig. 6.**
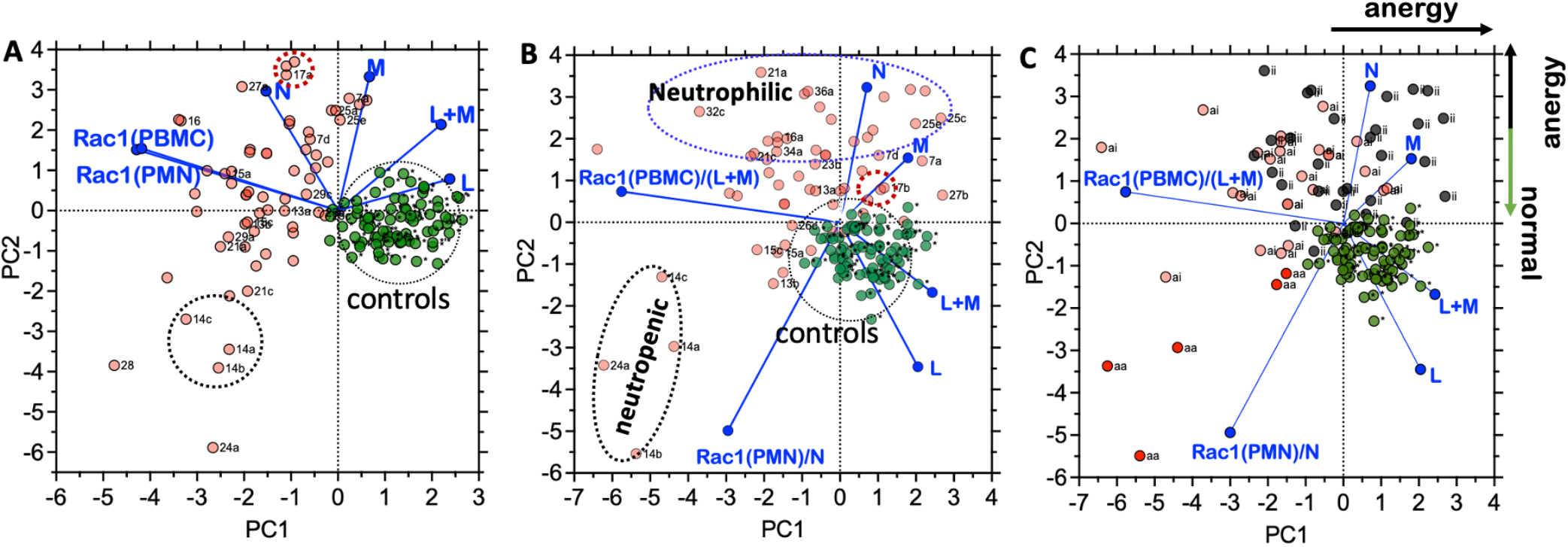
Principal Component Analysis (PCA) reveals different spatial distributions between uninfected control and blood culture-positive patients. **A.** Spatial distribution of infected patients (pink) relative to controls (green data points) The immune variables measured for each patient, Rac1 (PBMC), Rac1(PMN), L, L+M, M, and N, are denoted as blue PC loadings vectors. Rac1 (PBMC) and Rac1 (PMN) are measures of immune functionality of the total count of peripheral blood monocytes (PBMC) and polymorphonuclear cells (PMN), respectively. The magnitude of loadings vectors or data point coordinates indicates the weight of the variable and mutual relationships. Dotted circles around Patients #14 and #17 illustrate the opposite effects of neutropenia and neutrophilia when cell functionality is normalized to single cells in the following graph. **B.** PCA plot showing Rac1•GTP produced by singles for PBMCs: Rac1 (PBMC)/(L+M) and PMN: Rac1(PMN)/(N). Because of the high N count, Patient 17’s position in the PCA plot moves closer to controls, indicating diminished functionality of single cells in patients with neutrophilia. **C.** PCA plot showing GTP bound Rac1 normalized to cell count for PBMCs: Rac1 (PBMC)/(L+M) and PMN: Rac1(PMN)/(N). Plot identifies patients with the Rac1•GTP mediated response of PBMC and PMN (*aa*), active PBMC and inactive PMN (*ai*), and patients with a globally inactive immune response (*ii*). The original Biplot and PC scores figures were produced in GraphPad Prism version 9.3.1.

The Rac1(PBMC) and Rac1(PMN) vectors represent the level of functionality of the patient’s L, M, and N populations. The magnitude of the cosines of the angles between the vectors is related to mutual correlation, where small angles between two variables represent a strong correlation. Conversely, vectors mutually appearing at ~180° are negatively correlated, and those that seem nearly orthogonal (90°) are likely not correlated. In addition, proximity between *PC scores* and specific *PC loadings* vectors indicates a strong correlation. In **Fig. 6A,** Rap1(PBMC) and Rap1(PMN), represented by the group’s most extended vectors, present a strong correlation to each other along the PC1 negative axis. Control samples were grouped near the origin of PC1 and distributed on the negative axis overlapping the L vector. M and N vectors had more significant weight in PC2. The distribution of infected patient samples spanned both PC1 and PC2 axes.

When we examined the average GTPase activity of single cells, the Rac1(PMN) vector was redistributed to the negative PC2 on the opposite end of N and M. The coordinate shift by Rac1(PMN)/cell vector opposite to the N vector indicated a negative correlation between the two variables. This negative correlation between neutrophil counts and GTPase functionality highlighted a potential link between excessive neutrophilia and immune suppression linked to bacteremia. [51-53] Thus, patient samples clustering near the apex of the N vector expressed the cohort-peak neutrophil counts of low GTPase activity; e.g., the position of Patient #17 in the PCA shifts from a position of strong correlation to N and robust GTPase functionality in **Fig. 6A** to relatively weak Rac1(PMN)/cell functionality in **Fig. 6B**. Conversely, neutropenic patients (14 and 24) produced the highest levels of Rac1(PMN)/cell. This result indicated immune impairment due to limited cell counts, unrelated to functionality.

The aggregate responses from the infected and control patients’ leukocytes show the controls clustered near the origin and strongly correlated to L and L+M. The even clustering of controls between the PC1 and PC2 axis is consistent with a dynamic immune equilibrium[54] of the uninfected patients. However, this equilibrium is notably absent in a vast majority of the infected patients. To visualize the spatial distribution of patients based on immune functionality, we color-coded patient status depending on whether the Rac1(PBMC)/cell and Rac1(PMN)/cell data were both ≥ 3-fold greater than the controls (red, ***aa***), active *l*evel Rac1(PBMC) with *inactive* level Rac1(PMN) (pink, ***ai***) and *inactive* Rac1(PBMC) with Rac1(PMN) (dark, ***ii***). Most notably, the patients with high numbers of neutrophils (clusters next to N) appeared immune suppressed. Also, the immunosuppressed cluster cases were infected mostly by Klebsiella pneumoniae (KP), Streptococcus pneumoniae (SP), Staphylococcus aureus (SA), Pseudomonas aeruginosa (PA),[44] and Group B streptococcus (GBS)[55] are all pathogens known to thrive under conditions of immune compromise.

## Summary

Immune dysregulation in sepsis comprises the intersection of pro- and anti-inflammatory biomarkers [56]; thus, assays designed to test immune functionality will likely produce real-time actionable results.[57, 58] Proinflammatory cytokines are frequently cited as bacteremia biomarkers in the literature. [45, 56, 59-64] In our previous study[1] we showed how this interplay between circulating pro-and anti-inflammatory cytokines impacted the magnitude of the correlated GTPase signals. Thus, GTPase activity in patient leukocytes represents an integrated signaling outcome reflecting the composition of circulating cytokines due to infection. In this way, our assay delineated control patient samples, immune-active and immune-suppressed patients.

From a sepsis diagnostic perspective, a differential diagnosis of bacterial infection can be narrowed by the patient’s history, clinical examination, and *G-Trap immune functionality* test. Comorbidities such as various forms of cancer, cytotoxic chemotherapy, bone marrow, or tissue transplant are common causes of immunocompromise featuring neutropenia in the early stages, followed by depressed cell-mediated and humoral immunity. [65] Neutropenia increases patient susceptibility to opportunistic polymicrobial bacteremia *(e.g.,* pseudomonas aeroginosa, E. Coli (patient #14), and Klebsiella pneumoniae). Thus while patient #14’s neutrophil functionality was robust, the 100/ml neutrophil count elevated the patient’s susceptibility to infection based on a depleted cell count rather than functionality.[66] Establishing immune functionality in patients with comorbidities such as asplenic patients[65] lacking the ability to clear encapsulated bacteria are susceptible to opportunistic infections mainly caused by pneumococcus, and Haemophilus influenzae[66] would help streamline a differential diagnosis. T-cell immuno-suppression due to corticosteroids, cytokine antagonists, and monoclonals that deplete B-cells used to target connective tissue diseases, e.g., lupus erythematosus and rheumatoid arthritis, also promotes susceptibility to infection. Leukocyte functionality G-Trap tests could be an effective tool for effective diagnosis and narrow prescription of antibiotics based on local antibiograms.[66, 67]

There are limitations to this study that we will address in future work. First, our primary objective was to establish a proof-of-concept assay for interrogating the immune functionality of patients as a biomarker for the rapid assessment of presumptive bacteremia. Thus, our sample cohort was based on the availability of residual diagnostic samples with limited access to patient health records.[68, 69] Our pilot study lacked comprehensive IRB approval to access clinical patient variables that we could have used in aggregate with our current data. Thus, we were unable to characterize the trajectory of the disease and provide individualized insight into intervention strategy. Another limitation to be addressed in a future study is to characterize cytokine composition of the plasma, which can be used to crosscheck leukocyte functionality. Next, regarding our data quality, some variables have missing data. Specifically, the CBC with differential test was not always deemed necessary at the clinical level of our data set. Thus, such data are left blank. However, we established our present objective of using GTPase activity as a biomarker for delineating uninfected and infected patients. In summary, future blinded prospective studies will be necessary to show how host-immune functionality can complement existing methods in directing timely and accurate differential diagnosis.

## Materials and Methods

### Effector Proteins

GST-effector chimeras used for the studies are as follows: p21 activated kinase protein binding domain (PAK-1 PBD), a Rac1 effector, were obtained from Millipore Sigma. Rhotekin-RBD, a Rho effector protein, was purchased from Cytoskeleton (Denver, CO).

### Antibodies

Monoclonal Rac1 antibodies (Cat. # ARC03) were purchased from Cytoskeleton.

### Buffers

2X RIPA buffer: 100 mM Tris titrated with HCl to pH 7.4, 300 mM NaCl, 2 mM EDTA, 2 mM NaF, 2 mM Na_3_VO_4_, 2% NP-40, 0.5% sodium deoxycholate, and just before adding to the culture medium, 2 mM PMSF and 2X protease inhibitors. HHB buffer: 7.98 g/L HEPES (Na salt), 6.43 g/L NaCl, 0.75 g/L KCl, 0.095 g/L MgCl_2_ and 1.802 g/L glucose. HPSMT buffer, an intracellular mimic: 30 mM HEPES, pH 7.4, 140 mM KCl, 12 mM NaCl, 0.8 mM MgCl_2_, 0.01% Tween-20.

## Study design

The UNM HSC Institutional Review Board approved this study (UNM IRB#18-068). The study was divided into two patient populations. Our first goal was to collect residual diagnostic blood samples from patients admitted to UNM for a non-infectious disease cause. Residual peripheral blood test samples were collected from 120 subjects (50% female) presumed to be free of infection. Our second goal was to collect serial blood samples from the clinical lab after a positive confirmation of bacterial culture.

Serial blood samples ordered by a physician were collected daily and tested for bacteria culture growth, with excess samples stored at 4°C in the clinical laboratory till a bacteria culture was confirmed. In this way, serial blood test samples from 36 patients with a confirmed diagnosis of bloodstream infection were collected after standard of care (SOC) testing at TriCore Reference Laboratories (TRL). TRL provided the investigators with available complete blood counts with differential (CBC with diff.) for each subject. We did not include samples from 8 patients for lack of day 0-1 sample, limited availability, or sample loss.

### Isolation of Peripheral blood mononuclear cells (PBMCs) and Polymorphonuclear cells (PMNs) diagnostic samples

Whole blood samples were kept at 4 °C. Peripheral blood mononuclear cells (PBMC) consisted of a mixture of lymphocytes (T cells, B cells, and NK cells) and monocytes. Polymorphonuclear cells (PMNs), comprising primarily neutrophils with a small fraction of eosinophils, basophils, and mast cells, were isolated using a Ficoll-based density gradient. The layers containing the PBMC and PMNs were carefully recovered in 800 μl volumes, mixed with 800 μl PBS, centrifuged, and removed the excess solution, leaving the pellet in 25 μl volume.

### Functional assessment of PBMCs and PMNs with Rac1•GTP binding assays

The pelleted cells are lysed by adding 25 μl 2X RIPA at 4°C and centrifuged at 4°C in a cold room. 45 μl of cleared lysate was then analyzed for GTPase activity with G-Trap beads, functionalized with PAK-1 effector beads as previously described. [70] We used the CBC with diff. data to determine how much Rac1•GTP was produced by single cells.

### Statistical Analysis

We used GraphPad Prism 9.3.0 (GraphPad Software, La Jolla, CA) for statistical analysis. Principal component analyses and Pearson correlation coefficients were derived from the study of serial samples for each patient to visualize clustering between Rac-1 (PBMC), Rac1(PMN), lymphocytes (L), monocytes (M), and neutrophils (N).

## Acknowledgments

This research was funded by NIH National Center for Research Resources, the National Center for Advancing Translational Sciences: UL1TR001449, and the UNM Department of Pathology.

---

**Supplemental Table S1.** Raw Data Table of GTPase Activity and WBC blank spots indicate missing data. The Case ID letters refer to infecting pathogen, and (G±) is gram-positive or negative. (PM) Pasteurella multocida (G-); (SA) Staphylococcus aureus (G+), (MRSA) Methicillin-resistant Staphylococcus aureus (G+); (PA) Pseudomonas aeruginosa (G-); (SP) Streptococcus pyogenes (G+); (KP) Klebsiella pneumoniae (G-); (SL) Staphylococcus lugdunensis (G+); (PA) Pseudomonas aeruginosa (G-); EC Escherichia coli (G-); (GBS) Group B Streptococcus; (S) Salmonella (G-); (SC) Streptococcus constellatus (G+). * representative uninfected controls

| Case ID | Rac1(PBMC) | Rac1 (PMN) | WBC | L | N | M |
| --- | --- | --- | --- | --- | --- | --- |
| 9EC1 | 11120 | 13528.99 | 34 |  |  |  |
| 9EC2 | 7642.001 | 12486 | 7.98 |  |  |  |
| 10SA1 | 6437.998 | 3175. | 7.8 | 1.7 | 5.6 | 0.5 |
| 10SA2 | 10989 | 3454 | 6.86 |  |  | 0.6 |
| 11EC&SC-1 | 12673 | 2423 | 5.2 |  |  |  |
| 12GBS1 | 7708 | 6115 | 3.4 |  |  | 0.3 |
| 12GBS2 | 4960 | 2868 | 5.7 |  |  |  |
| 12GBS3 | 12873 | 3665.001 | 6.6 | 1.7 | 5.6 | 0.5 |
| 12GBS4 | 1853 | 1388 | 6.9 | 0.6 | 5.98 | 0.3 |
| 13EC1 | 5463.998 | 1488 | 8 | 1 | 6.7 | 0.4 |
| 13EC2 | 14974 | 11746 | 7.9 | 1.8 | 6.0 | 0.1 |
| 14EC1 | 6613.999 | 3187 | 0.9 | 0.7 | 0.5 | 0.1 |
| 14EC2 | 19178 | 7556.997 | 1 | 0.8 | 0.1 | 0.1 |
| 14EC3 | 14801.99 | 3349.999 | 1.8 | 0.5 | 1.4 | 0.1 |
| 15SA1 | 16317 | 16887 | 6.2 | 1 | 4.6 | 0.5 |
| 15SA2 | 12993 | 21886.01 | 5.2 | 1 | 3.4 | 0.6 |
| 15SA3 | 25700 | 2704 | 3.7 | 0.9 | 2.1 | 0.4 |
| 16PM1 | 50909.01 | 9788 | 19.4 | 0.5 | 17.1 | 0.9 |
| 16PM2 | 44474.96 | 12352 | 18.5 | 0.5 | 17.1 | 0.9 |
| 16PM3 | 60164.97 | 13008 | 17.4 | 0.5 | 17.1 | 1.4 |
| 17MRSA1 | 30180 | 9249 | 17.9 | 1.9 | 14.1 | 1.4 |
| 17MRSA2 | 37360 | 9815 | 18 | 2.1 | 15.1 | 1.5 |
| 6EA1 | 12682 | 5249 | 9.6 |  |  |  |
| 6EA2 | 9827 | 5026 | 11.6 |  |  |  |
| 7EC1 | 7270 | 1903 | 16.2 | 2.1 | 15.1 | 1.5 |
| 7EC2 | 2682 | 1504 | 13.4 | 0.7 | 11.9 | 1.3 |
| 7EC3 | 2291 | 1031 | 10.2 | 0.7 | 11.9 | 1.3 |
| 7EC4 | 1929 | 2567 | 9.9 | 0.7 | 10.9 | 1.5 |
| 8EC1 | 11241 | 2165 | 10.7 | 0.7 | 9 | 1 |
| 8EC2 | 11146 | 3679 | 7.4 | 0.8 | 5.5 | 0.5 |
| 8EC3 | 5030 | 5115 | 4.4 |  |  |  |
| 21PA1 | 881 | 801 | 17.4 | 0.2 | 17.4 | 0.5 |
| 21PA2 | 1062 | 872 | 10.8 | 0.3 | 9.5 | 1 |
| 21PA3 | 1006 | 854 | 8.1 | 0.4 | 7.4 | 0.2 |
| 23KP1 | 1491 | 1169 | 7.1 | 0.9 | 5.6 | 0.6 |
| 23KP2 | 1270 | 1028 | 9.7 | 0.7 | 8.4 | 0.4 |
| 23KP3 | 1257 | 988 | 7.1 | 0.9 | 5.6 | 0.6 |
| 24KP1* | 1306 | 1101 | 0.2 | 0.2 | 0.1 | 0.1 |
| 24KP2* | 790. | 831 | 0.4 | 0.3 | 0.1 |  |
| 22SL1 | 1062 | 6218 | 10.2 | 0.6 | 8.9 | 0.6 |
| 22SL-2 | 1143 | 983 |  |  |  |  |
| 22SL-3 | 940 | 2581 |  |  |  |  |
| 22SL-4 | 955 | 1000 |  |  |  |  |
| 25SA1 | 1025 | 1867 | 47.7 | 1 | 44.8 | 1.4 |
| 25SA2 | 8486 | 3084 | 40.3 | 0.8 | 37.1 | 2.4 |
| 25SA3 | 1976 | 1379 | 33.8 | 1.7 | 30.8 | 1.4 |
| 25SA4 | 3839 | 1145 | 32 | 1 | 30.1 | 1 |
| 25SA5 | 2338 | 1487 | 28.2 | 1.4 | 25.4 | 1.2 |
| 26SP1 | 4716 | 1652 | 8.1 | 0.7 | 6.7 | 0.5 |
| 26SP2 | 1588 | 4145 | 5.1 | 1 | 3.6 | 0.4 |
| 26SP3 | 1547 | 1399 | 4.6 | 0.8 | 3.4 | 0.3 |
| 27SP1 | 11295 | 12714 | 28 | 0.8 | 25.5 | 1.4 |
| 27SP2 | 6320 | 3691 | 29.5 | 4.4 | 24.5 | 0.6 |
| 27SP3 | 6733 | 3463. | 15.7 | 2.4 | 12.9 | 0.3 |
| 28EC1 | 4532 | 1105 | 4.1 | 0.1 | 3.6 | 0.1 |
| 28EC2 | 3406 | 3576 | 15.4 |  |  |  |
| 28EC3 | 2496 | 4609 | 10.3 |  |  |  |
| 28EC4 | 7905 | 2392 | 5.6 |  |  |  |
| 28EC5 | 3173. | 2313 | 4.2 |  |  |  |
| 28EC6 | 3904. | 1501 |  |  |  |  |
| 29SP1 | 2880 | 2230 | 18.9 | 0.6 | 18.1 | 0.2 |
| 29SP3 | 3362. | 1882 | 11.6 | 0.7 | 10.1 | 0.2 |
| 29SP4 | 1918 | 1245 |  |  |  |  |
| 29SP5 | 5945. | 1569 | 8.1 | 1.3 | 5.8 | 0.5 |
| 30SA1 | 9476. | 2934 | 10.7 | 0.5 | 9.2 | 1.1 |
| 30SA2 | 8895 | 3594 | 11.6 | 0.9 | 9.2 | 1.5 |
| 31SA1 | 3002. | 6002. | 0.1 | 0.1 |  |  |
| 31EC2 | 4281. | 3009. |  |  |  |  |
| 31EC3 | 2568 | 8990. |  |  |  |  |
| 31EC4 | 1622 | 3856. | 0.19 |  |  |  |
| 31EC5 | 4102. | 1017 |  |  |  |  |
| 31EC6 | 1885 | 10006 | 0.1 |  |  |  |
| 32SA1 | 4216. | 1203 | 18.6 | 0.8 | 17.6 | 0.2 |
| 32SA2 | 1833 | 2366 | 24.3 | 0.3 | 23.3 | 0.7 |
| 32SA3 | 2346 | 1887 | 14.3 | 0.2 | 13.9 | 0.2 |
| 32SA4 | 2415 | 902 | 10.2 | 0.6 | 8.8 | 0.2 |
| 32SA5 | 2154 | 997. |  |  |  |  |
| 33SA | 4670. | 1780 | 8.6 | 1.4 | 6.2 | 0.9 |
| 34SP1 | 10787 | 5908. | 15.3 | 0.5 | 13 | 0.6 |
| 34SP2 | 7977. | 5881. | 14.1 |  |  |  |
| 34SP3 | 10038 | 4835. | 12.6 |  |  |  |
| 34SP4 | 8227. | 5179 | 11.7 |  |  |  |
| 34SP5 | 9071 | 6561. | 10.8 |  |  |  |
| 35EC1 | 4974. | 5927 | 11.2 | 2 | 7.8 | 1.4 |
| 35EC3 | 4728. | 3412 | 8 |  |  |  |
| 35EC4 | 3372 | 2518 | 13.2 |  |  |  |
| 36SP1 | 3164 | 1236 | 26.9 | 0.5 | 25.6 | 0.5 |
| 36SP2 | 5972. | 2133 | 11.9 |  |  |  |
| 36SP3 | 4261. | 1779 |  |  |  |  |
| 36SP4 | 3329. | 2267 |  |  |  |  |
| * | 2459 | 3055 | 7.2 | 2.8 | 3.4 | 0.7 |
| * | 3750 | 3297 | 6 | 1 | 4.1 | 0.7 |
| * | 3669 | 3727 | 6.9 | 1.9 | 6.1 | 0.8 |
| * | 2473 | 2159 | 4.7 | 2.2 | 2.3 | 0.7 |
| * | 2203 | 2073 | 6.1 | 2.4 | 3 | 0.5 |
| * | 7999 | 2373 | 5.3 | 2.1 | 2.5 | 0.4 |
| * | 2290 | 3077 | 5.9 | 1.2 | 4 | 0.6 |
| * | 3160 | 3440 | 5.8 | 1.8 | 3.4 | 0.4 |
| * | 5272 | 3762 | 7 | 2.7 | 3.7 | 0.5 |
| * | 2496 | 1234 | 6.8 | 2.2 | 3.9 | 0.5 |
| * | 5846 | 1431 | 6.3 | 1.4 | 4.2 | 0.3 |
| * | 2293 | 1353 | 4.9 | 1.6 | 2.7 | 0.4 |
| * | 2023 | 1241 | 6.8 | 2.5 | 3.3 | 0.8 |
| * | 1660 | 1145 | 4.4 | 1.5 | 2.4 | 0.4 |
| * | 2570 | 1518 | 5 | 1.2 | 3.1 | 0.5 |
| * | 2731 | 1214 | 6.4 | 2.1 | 3.4 | 0.6 |
| * | 1927 | 964 | 5.2 | 2.5 | 2.1 | 0.5 |
| * | 3074 | 1395 | 5.7 | 2.4 | 2.9 | 0.3 |
| * | 2295 | 1536 | 4.8 | 1.4 | 2.9 | 0.4 |
| * | 2315 | 1391 | 6.1 | 2 | 3.6 | 0.4 |
| * | 1804 | 1071 | 5.3 | 1.7 | 3.0 | 0.5 |

